# Surface *N*-acetylglucosamine dynamics in bovine spermatozoa: from epididymal transit to oviductal epithelial cell binding

**DOI:** 10.64898/2026.08.07.743513

**Authors:** Pablo Ariel Alvarez, Natalia Lorena Leiva, Lorena Carvelli, Inmaculada Robina, Miguel Angel Sosa, Andrea Carolina Aguilera

## Abstract

In mammals, the sperm surface glycocalyx undergoes extensive remodeling during epididymal maturation, which is required for spermatozoa to acquire the ability to reach and bind the oviductal epithelium. N-acetylglucosamine (GlcNAc)-containing glycans are dynamically modified during bovine epididymal maturation, but whether these changes are regulated by specific epididymal enzymes, and whether surface GlcNAc contributes to sperm–oviduct epithelial interactions, remain poorly defined. Here, we addressed this gap in bovine spermatozoa by examining how surface GlcNAc changes during sperm maturation and functional activation—from epididymal maturation through capacitation and the acrosome reaction—and by asking whether these changes relate to the ability of spermatozoa to bind the bovine oviductal epithelium. Surface GlcNAc, assessed by WGA reactivity, changed progressively from the proximal caput to the distal cauda epididymidis, shifting toward a more homogeneous population; this progression was accompanied by a shift in GlcNAc-bearing protein profiles, while GlcNAc remained predominantly localized to the acrosomal region throughout maturation. Building on our previous finding that β-*N*-acetylglucosaminidase (β-NAG) content increases significantly in the cauda epididymis of reproductively mature bulls, we hypothesized that this regional increase in enzymatic activity underlies the observed changes in surface GlcNAc. Consistent with this hypothesis, incubation of caput spermatozoa with cauda epididymal fluid reduced WGA labeling, an effect blocked by the selective β-NAG inhibitor VP150, identifying luminal β-NAG as an active contributor to GlcNAc remodeling during epididymal transit. In ejaculated spermatozoa, capacitation induced only minor changes in surface GlcNAc, whereas the calcium ionophore-induced acrosome reaction produced a marked reduction in WGA reactivity and acrosomal labeling, consistent with glycoprotein loss during acrosomal exocytosis. Functionally, spermatozoa that bound to bovine oviductal epithelial cell (BOEC) monolayers were preferentially WGA-positive, and pre-incubation of spermatozoa with WGA significantly reduced sperm adhesion, supporting a role for surface GlcNAc in sperm–oviduct epithelial interactions. Together, these findings show that surface GlcNAc is dynamically regulated during sperm maturation and functional activation, with progressive changes during epididymal transit that are partly mediated by luminal β-NAG activity, followed by redistribution during capacitation and acrosomal exocytosis. The association between surface GlcNAc and sperm–BOEC adhesion further supports a role for GlcNAc-containing glycans in sperm–oviduct epithelial interactions, providing a potential mechanistic framework for glycocalyx-mediated sperm selection in cattle.

## 1. INTRODUCTION

Fertilization in mammals is the result of coordinated molecular and cellular events that begin long before gamete fusion. The dynamic remodeling of the sperm surface glycocalyx, a dense carbohydrate-rich coat that surrounds spermatozoa and mediates interactions throughout the male and female reproductive tracts, and this remodeling is critical to the fertilization process [1,2]. In mammals, glycosylation is a key post-translational modification in male reproduction, playing roles in spermatogenesis, sperm maturation, immune protection, cell-cell recognition, and fertilization [2,3]. Among sperm surface glycans, GlcNAc-containing structures are particularly important, and they become evident during epididymal maturation and capacitation, contributing to sperm-oviduct and sperm-zona pellucida interactions [4,5].

Sperm maturation during epididymal transit is a highly regulated post-testicular process through which male germ cells acquire progressive motility and fertilizing competence. Although spermatozoa are transcriptionally and translationally silent, their surface proteins undergo substantial remodeling through addition, removal, or modification of glycan structures via interactions with the epididymal luminal environment [1,6]. The epididymis has been aptly described as an “extracellular Golgi” given its capacity to reshape the sperm glycocalyx through secreted enzymes, luminal proteins, and membrane-derived vesicles [2,7]. Among the enzymatic actors present in epididymal fluid, glycan-modifying enzymes, including glycosidases and glycosyltransferases, are thought to drive the stage-specific remodeling of surface glycoconjugates that accompanies sperm maturation [8,9]. In particular, β-*N*-acetylglucosaminidase (β-NAG) (EC 3.2.1.52), which hydrolyzes terminal GlcNAc residues from glycoconjugates, has been detected in the epididymal fluid of several species and is a plausible regulator of sperm surface GlcNAc content. Previous work from our group demonstrated significantly elevated β-NAG activity in the cauda epididymal fluid of sexually mature bulls, suggesting a role for this glycosidase in sperm surface remodeling during epididymal transit [10]. Recent evidence further underscores the physiological importance of GlcNAc-related pathways in sperm biology: the nucleotide sugar transporter SLC35G3 has been identified as essential for GlcNAc incorporation into sperm glycoproteins, since its disruption results in male infertility in mice [11], and lectin-based glycan profiling has linked differential GlcNAc abundance to fertilizing outcomes in buffalo bulls [12].

Upon ejaculation and entry into the female reproductive tract, spermatozoa interact with the oviductal epithelium to establish a sperm reservoir, a specialized microenvironment that prolongs sperm lifespan, maintains fertilizing competence, and regulates the timing of capacitation until ovulation occurs [13]. All these interactions are largely mediated by carbohydrate–lectin recognition mechanisms at the sperm–epithelial interface, involving glycoconjugates expressed on both gametes and oviductal epithelial cells [14,15]. Several sperm surface glycan structures, including L-fucose-, D-mannose, and sialic acid-containing moieties, have been associated with bovine sperm binding to oviductal epithelial cells [14,16]. Moreover, GlcNAc-containing glycans and their binding partners are attractive candidates for mediating these adhesive events. However, although surface carbohydrate remodeling has been described in bovine spermatozoa [8,17–19], a causal, enzyme-specific link between epididymal glycosidase activity and GlcNAc remodeling has not been established, and direct functional evidence for a role of GlcNAc in sperm–bovine oviductal epithelial cell (BOEC) adhesion is lacking.

It is well known that capacitation and the acrosomal reaction are accompanied by substantial remodeling of the sperm glycocalyx, including alterations in the distribution and relative abundance of specific glycan residues [20,21]. Using fluorescently conjugated lectins, including Wheat Germ Agglutinin (WGA), which binds with high affinity to terminal GlcNAc and sialic acid residues, changes in sperm surface carbohydrate profiles during these functional transitions in several species have been documented [18,19]. However, whether specific epididymal enzymes actively drive this GlcNAc remodeling during maturation, and whether the resulting changes in surface GlcNAc are functionally relevant for sperm binding to the oviductal epithelium, has not been addressed.

In this context, we hypothesized that surface GlcNAc is progressively modifed during epididymal transit with the participation of luminal β-NAG activity, is redistributed during acrosomal exocytosis, and functionally mediates sperm binding to the oviductal epithelium. To test this hypothesis, the present study was designed to provide a mechanistic and functional characterization of surface GlcNAc dynamics in bovine spermatozoa throughout their journey from the epididymis to the oviduct. Rather than solely documenting descriptive changes in GlcNAc abundance, we specifically sought to identify the epididymal enzymatic activity responsible for this remodeling and to establish its functional consequence for sperm–oviductal epithelial cell (BOEC) adhesion. To this end, we first characterized changes in surface GlcNAc content and glycoprotein profile during epididymal transit from caput to cauda and then evaluated whether luminal β-NAG activity directly mediates GlcNAc remodeling on immature spermatozoa, providing a candidate enzymatic driver for the changes observed *in vivo*. We further examined the dynamics of GlcNAc redistribution during *in vitro* capacitation and the acrosome reaction, and, critically, assessed the association between surface GlcNAc exposure and the ability of ejaculated spermatozoa to bind to BOEC monolayers. Collectively, these findings move beyond a descriptive account of sperm glycocalyx remodeling to identify a specific epididymal enzyme driving GlcNAc dynamics and to demonstrate its functional link to sperm–oviduct epithelial recognition in cattle.

## 2. MATERIALS AND METHODS

### 2.1. Antibodies and Reagents

FITC-conjugated wheat germ agglutinin (WGA; cat. no. L4895) and biotin-conjugated WGA (cat. no. L5141) were purchased from Sigma-Aldrich (St. Louis, MO, USA). The fluorogenic substrate 4-methylumbelliferyl *N*-acetyl-β-D-glucosaminide (cat. no. M2133), used for β-*N*-acetylglucosaminidase activity assays, was also obtained from Sigma-Aldrich. Nitrocellulose membranes (0.45 μm pore size) were purchased from GE Healthcare (Freiburg, Germany). Calcium ionophore A23187 (cat. no. C7522), heparin (cat. no. H3149), HRP-conjugated streptavidin (cat. no. 189733), anti-β-tubulin antibody (cat. no. T8328), anti-vimentin antibody (cat. no. V6630), FITC-conjugated goat anti-rabbit IgG secondary antibody (cat. no. F6005), and Hoechst 33342 (cat. no. B226) were purchased from Sigma-Aldrich. The anti-cytokeratin antibody (cat. no. ab9377) was purchased from Abcam (Cambridge, UK) and UltraCruz™ Fluorescence Mounting Medium (cat. no. sc-2494) was acquired from Santa Cruz Biotechnology.

The enhanced chemiluminescence (ECL) reagent was prepared in-house by combining 1.25 mM luminol and 198 μM *p*-coumaric acid in 100 mM Tris-HCl (pH 8.5), as previously described [22]. Unless otherwise stated, all other reagents were of analytical grade and purchased from Sigma-Aldrich.

### 2.2. Animals and biological samples

All procedures were performed as previously described by Aguilera et al. (2017) [8]. No animals were sacrificed specifically for the purpose of this study. All bovine reproductive tracts were obtained as by-products from a local commercial slaughterhouse (María del Carmen, Corralitos, Mendoza) in strict accordance with local regulations for the handling and processing of animal subproducts. A total of 12 sexually mature *Aberdeen Angus* bulls (18-24 months old) of proven fertility were used as the source of biological material. For epididymal studies, 7 bulls were sampled post-mortem at the slaughterhouse, and both epididymides (n=14 organs) were collected from each animal for processing. For experiments involving ejaculated spermatozoa, frozen-thawed straws from reproductively mature Aberdeen Angus bulls (2–5 years of age) were purchased and utilized as independent biological replicates. Each epididymis was carefully dissected, and the three main segments (caput, corpus, and cauda) were processed separately. Tissues from each region were minced with a stainless-steel blade and suspended at 1:3 (w/v) in Hank’s solution at 37 °C for 30 min with gentle manual agitation. After settling for 10 min, the supernatant containing spermatozoa and luminal fluid was collected and centrifuged at 400 × g for 5 min to separate spermatozoa from the fluid. For experiments involving flow cytometry evaluation, cauda epididymal luminal content was collected by retrograde flushing with Hank’s solution and then spermatozoa and fluid were then separated as described above. Spermatozoa from the caput were obtained from small incisions made on selected tubules and applying slight pressure to the proximal region. Only intraluminal samples free of blood or tissue debris were further processed. Gametes were washed three times with Hank’s solution and finally pelleted at 1,000 × g for 10 min.

### 2.3. Obtention of ejaculated spermatozoa, capacitation and acrosome reaction induction

Cryopreserved semen straws from reproductively mature *Aberdeen Angus* bulls (approximately 30 × 10 spermatozoa/straw) were thawed at 37 °C for 30 s and immediately diluted in pre-warmed non-capacitating TALP medium (TALP-NC). Motile spermatozoa were then selected by swim-up in either capacitating TALP (TALP-C) or non-capacitating TALP (TALP-NC), prepared according to Parrish et al. (1988) (23). TALP-C was prepared from the TALP-NC supplemented with heparin (10 µg/mL) and BSA (6 mg/mL). Spermatozoa recovered immediately after thawing, prior to any further incubation, were designated as the “freshly thawed condition” (Ej). Motility and viability were assessed by light microscopy (×400 magnification) before use in subsequent trials (capacitation, acrosome reaction, or BOEC-binding). For capacitation, spermatozoa were incubated in TALP-C for 4 h at 38.5 °C in a humidified atmosphere of 5% CO. To induce the acrosome reaction, capacitated spermatozoa were subsequently exposed to 15 µM calcium ionophore A23187 for 15 min under the same atmospheric conditions.

### 2.4. Detection of GlcNAc in nitrocellulose-immobilized sperm glycoproteins

GlcNAc-containing glycoproteins were detected in sperm protein extracts by lectin blot using biotinylated wheat germ agglutinin (WGA-biotin). The protein extraction strategy varied according to the experimental purpose: enriched membrane fractions were used for epididymal maturation studies, whereas total protein extracts were used for capacitation and acrosome-reaction studies (rationale detailed in the Discussion). Protein concentrations were determined using the Lowry colorimetric method [24]. For lectin blot analysis, either 45 µg of membrane protein extracts or sperm total protein from 3 × 10 spermatozoa were analyzed on 10% SDS–PAGE gels under non-reducing conditions. Then, the proteins were electrotransferred onto nitrocellulose membranes (0.45 µm pore size) for 90 min at 250 V. Non-specific sites on nitrocellulose were then blocked for 1 h with 6% skim milk prepared in PBS containing 0.05% Tween-20 (PBS-T), washed three times with PBS-T, and incubated overnight (ON) with biotin-conjugated WGA (5 µg/mL in PBS-T), at 4°C. After extensive washing, membranes were incubated for 1 h with HRP-conjugated streptavidin (1:10000 dilution in PBS-T). Lectin-reactive bands were visualized using an enhanced chemiluminescence system according to Haan and Behrmann [22], and images were acquired using an ImageQuant LAS 4000 imaging system. Band intensities were quantified by densitometric analysis. Ponceau S staining or β-tubulin immunodetection was used as loading control, as appropriate for each experiment.

### 2.5. Fluorescence microscopy

The distribution of GlcNAc residues on the sperm surface was assessed using a fluorescence-based lectin-labeling assay adapted from Aguilera et al. (2017) [8]. Spermatozoa retrieved from the caput, corpus and cauda epididymis were resuspended in PBS supplemented with 0.1% polyvinylpyrrolidone (PBS/PVP) and fixed in 2% paraformaldehyde (PAF) for 10 min at room temperature. Following fixation, cells were seeded onto glass slides previously coated with 10 mM poly-L-lysine for 30 min to promote adhesion. Slides were blocked for 1 h at room temperature with 5% horse serum prepared in PBS/PVP and subsequently incubated with FITC-conjugated wheat germ agglutinin (WGA–FITC, 10 µg/mL in PBS/PVP containing 1% horse serum). Samples were mounted and examined under a Nikon Eclipse 80 fluorescence microscope (Nikon, Japan). Images were acquired using identical exposure settings for all experimental conditions.

### 2.6. Flow cytometry

Sperm surface-exposed *N*-acetylglucosamine (GlcNAc) residues were quantified by flow cytometry as previously described [8], with modifications adapted to the present study. Epididymal and ejaculated spermatozoa were washed three times in Hank’s balanced salt solution and centrifuged at 1,000 × g for 10 min. Cells were subsequently fixed in 2% paraformaldehyde (PAF) prepared in PBS/PVP for 10 min at room temperature, blocked for 1 h at 37 °C with PBS/PVP containing 5% horse serum, and washed three times with PBS/PVP prior to lectin incubation. For each condition, 5 × 10 spermatozoa were incubated in the dark with FITC-conjugated wheat germ agglutinin (WGA-FITC; 5 µg/mL) in a final volume of 100 µL for 2 h at room temperature. Specificity of lectin binding was verified by incubation in the presence of 0.5 mM N-acetyl-D-glucosamine (GlcNAc), which competitively inhibited WGA binding. After staining, cells were washed twice with PBS/PVP and analyzed using a BD FACSAria III flow cytometer (BD Biosciences, USA) equipped with FACSDiva software. Sperm populations were identified by establishing gates on forward scatter (FSC) and side scatter (SSC) dot plots based on their characteristic size and granularity parameters. FITC fluorescence was excited at 488 nm and collected using a 525 nm band-pass filter. A minimum of 10,000 events was counted for each sample. The percentage of WGA-positive spermatozoa and the mean fluorescence intensity (MFI) were quantified using FlowJo software version 10.1.

### 2.7. Enzymatic activity assays

The activity of β-*N*-acetylglucosaminidase (β-NAG) was determined using 4-methylumbelliferyl *N*-acetyl-β-D-glucosaminide as a fluorogenic substrate, following procedures from our laboratory [8]. One unit of enzymatic activity was defined as the amount of enzyme required to release 1 nmol of 4-methylumbelliferone (4-MU) per minute under the assay conditions. Reactions were conducted in 0.2 mM phosphate buffer (pH 6.5) at 37 °C for the specified incubation period. The reactions were stopped with 0.2 M sodium carbonate, and the fluorescence of 4-methylumbelliferone was measured using a microplate spectrofluorometer (excitation/emission: 365/450 nm). To normalize enzymatic activity (as specific activity) protein concentration was measured in each sample using the Lowry method.[24]

### 2.8. Isolation and culture of bovine oviductal epithelial cells (BOECs)

Bovine oviducts were collected from sexually mature *Aberdeen Angus* cows (18-20 months old) from a local slaughterhouse and handled following the procedures described by Lamy et al. [25]. Briefly, reproductive tracts were transported to the laboratory at 37 °C and processed within 2 h post-mortem. Based on ovarian morphology and the presence or absence of a functional corpus luteum, as described by Ireland et al. [26], only those oviducts at the peri-ovulatory stage were included in the study. Upon arrival, both oviducts from each animal were dissected and maintained free of surrounding connective tissue. To isolate the epithelium, each oviduct was first trimmed of surrounding connective tissue and rinsed with warm TCM-199. The oviductal epithelial cells (BOECs) were recovered by applying gentle external pressure along the entire length of the oviduct using a sterile glass slide. To remove debris and contaminating cells the collected mucosal material was washed three times with HEPES-buffered TCM-199 by sedimentation for 10 min each. The resulting pellet was resuspended in TCM-199, supplemented with 20% heat-inactivated fetal calf serum and Penicillin–Streptomycin (100 U/mL/100 µg/mL). BOECs were then seeded into 96 well plate (Nunc, Roskilde, Denmark) and cultured in a humidified incubator at 38.8 °C with 5% CO. The culture medium was replaced after 48 h, and thereafter half-renewed every 48 h until the cells reached 90-100% confluence (typically 5–6 days). Confluent monolayers were used for subsequent experiments as described below.

### 2.9. Characterization of bovine oviductal epithelial cells (BOECs)

The epithelial phenotype of confluent BOEC monolayers was confirmed by immunofluorescence using cytokeratin and vimentin as markers. For immunofluorescence analysis, BOEC monolayers were fixed with 2% paraformaldehyde, permeabilized with 0.1% Triton X-100, and blocked with 5% bovine serum albumin (BSA) in PBS. Cells were then incubated with anti-cytokeratin antibody (1:200 dilution) or anti-vimentin antibody (1:100 dilution) followed by a FITC-conjugated anti-rabbit secondary antibody (1:50 dilution). Nuclei were counterstained with Hoechst 33342, and samples were examined under fluorescence microscopy. The proportions of cytokeratin-positive (Cyt) and vimentin-positive (Vim) populations were quantified.

### 2.10. Co-incubation of sperm with BOECs

Frozen–thawed bovine ejaculated spermatozoa were used in all co-culture assays. Straws (0.25 mL; approximately 30 × 10 spermatozoa per straw) were thawed in a water bath at 37 °C for 1 min and washed in non-capacitating Tyrode’s medium (TALP-NC) supplemented with 200 nM sodium pyruvate, 10 mM lactate and antibiotics (100 IU/mL penicillin and 100 µg/mL streptomycin). After centrifugation, the sperm pellet was resuspended in TALP-NC and sperm concentration was determined by counting in a Neubauer chamber. BOECs monolayers were used at full confluence. Before co-incubation, BOECs were washed twice with TALP-NC. Spermatozoa were then added at a final concentration of 4 × 10 cells/mL. Co-cultures were maintained in TALP-NC at 38.8 °C in a humidified atmosphere with 5% CO for 20 min.

### 2.11. Quantification of spermatozoa bound to BOECs

At the end of co-incubation, BOECs monolayers were washed vigorously three times with TALP-NC to remove unbound and weakly attached spermatozoa. For quantification under standard conditions, cells were fixed with 2% paraformaldehyde (PAF) in PBS for 20 min at room temperature, washed in PBS, and stained with Hoechst 33342 to visualize nuclei. In experiments assessing membrane integrity, spermatozoa were stained immediately after co-culture with propidium iodide (PI) and imaged live without fixation. Samples were observed using a Zeiss AxioObserver Z1 fluorescence microscope (Zeiss, Oberkochen, Germany) equipped with a spectral imaging detector. Excitation wavelengths of 405 nm and 633 nm were used for Hoechst and PI, respectively, using 10× or 20× (NA 0.5 objectives). For each experimental condition, 10 randomly selected fields (0.18 mm² each) were imaged. The number of BOECs and the spermatozoa attached to their surface was quantified using Fiji/ImageJ software. Particle analysis was performed by setting size thresholds corresponding to the dimensions of sperm heads and epithelial cells, followed by manual verification to confirm correct discrimination between both populations. The results were expressed as the number of spermatozoa bound per 100 BOECs.

### 2.12. Statistical analyses

All data sets were first assessed for normality and homogeneity of variance. Depending on the distribution, comparisons between two groups were performed using an unpaired *t*-test, while comparisons among more than two groups were analyzed by one-way ANOVA using the GraphPad Prism software, version 8. When ANOVA indicated significant differences, either Dunnett’s test or Tukey–Kramer’s multiple comparisons test was applied, as appropriate for each experiment. Statistical significance was considered at p < 0.05. Results are expressed as mean ± SEM unless otherwise indicated.

### Experimental design

#### Experiment 1. Qualitative and quantitative assessment of GlcNAc residues in caput, corpus and cauda epididymal spermatozoa

This experiment was designed to evaluate the distribution and content of surface-exposed GlcNAc residues in epididymal spermatozoa (from caput, corpus, and cauda epididymis). Epididymal spermatozoa were analysed by flow cytometry and fluorescence microscopy using FITC-conjugated WGA, following procedures adapted from Aguilera et al. 2017 [8]. Relative fluorescence intensity and the proportion of WGA-positive sperm subpopulations were quantified to assess changes in carbohydrate content occurring during epididymal transit. To further characterize molecular remodelling associated with sperm maturation, GlcNAc-bearing glycoproteins were analysed by lectin blot. For this purpose, enriched membrane fractions were isolated from epididymal spermatozoa using hypo-osmotic extraction protocol adapted from Jeyendran et al. 1984 [27], and previously implemented in our laboratory [8]. Briefly, spermatozoa (≤1 × 10 cells/mL) were incubated for 2 h at 37 °C in a hypo-osmotic solution containing 25 mM sodium citrate dihydrate and 75 mM fructose. Samples were subsequently sonicated on ice until sperm tail detachment (confirmed microscopically) and centrifuged at 4000 × g for 15 min to eliminate intact cells. The resulting supernatants were further centrifuged at 15000 × g for 30 min to obtain enriched membrane fractions. Pellets were resuspended in 0.05 M Tris–HCl buffer (pH 7.2) containing 0.5% saponin, 50 mM EDTA, and 1 mM PMSF before lectin blot analysis as described above. Ponceau S staining was used as loading control for epididymal samples.

#### Experiment 2. Effect of cauda epididymal fluid on the GlcNAc content of caput spermatozoa

Based on previous findings from our group demonstrating increased β-*N*-acetylglucosaminidase (β-NAG) activity in the epididymal cauda luminal fluid of sexually mature bulls [10], we investigated whether soluble factors present in the cauda luminal fluid contribute to the remodeling of GlcNAc residues on immature spermatozoa. Freshly isolated caput spermatozoa were incubated with crude cauda epididymal fluid under physiological conditions, as previously described [8]. Following incubation, the abundance of surface-exposed GlcNAc residues was assessed by flow cytometry using FITC-conjugated wheat germ agglutinin (WGA-FITC). To evaluate the specific involvement of β-NAG activity in this process, parallel incubations were performed in the presence of the selective β-NAG inhibitor VP150 (1 mM), provided by the Department of Organic Chemistry, Faculty of Chemistry, University of Seville. [28]. The inhibitor was added to the cauda fluid and pre-incubated for 1h before its addition to sperm samples. Controls were performed with sperm treated with the vehicle alone (9:1, H O:MeOH). The molecular structure of VP150 is shown in Supplementary Fig. 1A. After incubation, spermatozoa were washed and processed for flow cytometric analysis as described above.

#### Experiment 3. Changes in GlcNAc content in ejaculated spermatozoa under different conditions: capacitated, or acrosome-reacted spermatozoa

This experiment aimed to determine whether sperm functional status is related to the abundance and distribution of GlcNAc residues on the gamete surface. Frozen–thawed ejaculated spermatozoa were analysed after subjecting them to different conditions: after incubation in non-capacitating medium (NCp), after incubation in capacitating medium (Cp), and following induction of the acrosome reaction using the calcium ionophore A23187 (RA). They were all compared to those sperm immediately after thawing (Ej). For each condition, spermatozoa were incubated with FITC-conjugated WGA and analysed by flow cytometry to quantify changes in GlcNAc-associated fluorescence. Fluorescence microscopy was additionally performed to evaluate the spatial distribution of lectin labelling under each functional state. To investigate changes in the patterns of GlcNAc-bearing proteins associated to sperm activation and acrosomal exocytosis, total protein extracts were prepared from spermatozoa from each experimental condition. In this case, total protein extracts were intentionally used to preserve the complete spectrum of proteins potentially affected during capacitation and the acrosome reaction. This strategy was considered particularly relevant given that acrosomal exocytosis involves extensive membrane fusion events and release of acrosomal components, which could substantially modify lectin-binding patterns. Protein extracts were separated by SDS–PAGE, electrotransferred onto nitrocellulose membranes and analysed by lectin blot using biotinylated WGA as described above. Lectin-reactive bands were visualized using HRP-conjugated streptavidin and enhanced chemiluminescent detection. β-tubulin immunodetection was used as loading control for ejaculated sperm protein extracts.

#### Experiment 4. Association between surface GlcNAc residues and sperm binding to BOECs

This experiment was designed to evaluate whether the presence of surface-exposed GlcNAc residues is associated with the ability of ejaculated spermatozoa to interact with bovine oviductal epithelial cells (BOECs). Ejaculated spermatozoa were co-incubated with BOEC monolayers under the conditions described above. Following co-culture, bound spermatozoa were identified by fluorescence microscopy after staining with FITC-conjugated wheat germ agglutinin (WGA-FITC). To further characterize the non-adherent population, the supernatant containing unbound spermatozoa was collected and analysed by flow cytometry. Cells were double-stained with WGA-FITC and propidium iodide (PI), allowing discrimination of subpopulations based on GlcNAc exposure and membrane integrity (e.g., WGA PI and WGA PI). To assess the functional involvement of GlcNAc residues in sperm–BOEC interaction, spermatozoa were pre-incubated with unconjugated WGA (10 µg/mL) for 30 min prior to co-incubation with BOEC monolayers. Following treatment, sperm binding was quantified as described above.

## 3. RESULTS

### 3.1. The surface content of *N*-acetylglucosamine (GlcNAc) changes during epididymal transit

To determine whether the presence of GlcNAc residues on the sperm surface varies along the bovine epididymis, spermatozoa recovered from the caput, corpus, and cauda were analyzed using WGA-FITC. Flow cytometry revealed two distinct populations with low (WGA^+^) and high (WGA^++^) WGA staining. In caput and corpus spermatozoa, these two populations coexisted, indicating a heterogeneous distribution of surface GlcNAc within these segments. Toward the cauda, this bimodal pattern was largely resolved, with the majority of spermatozoa displaying uniform WGA staining, indicating that the sperm population becomes markedly more homogeneous with respect to surface GlcNAc as epididymal transit proceeds. Among WGA^++^ spermatozoa, the RFI also varied in FITC intensity, and significant differences in RFI were detected among the three epididymal segments (P < 0.05), with a progressive increase in the WGA^++^ population during epididymal transit (Fig. 1A–B). By fluorescence microscopy, no major changes in GlcNAc distribution were observed during epididymal maturation, as the sugar residue remained mostly localized to the acrosomal region of spermatozoa across all three segments (Fig. 1C). Lectin blot analysis further showed that this shift toward a more homogeneous surface phenotype was paralleled by a simplified GlcNAc-bearing protein pattern in cauda membrane fractions, with the disappearance of several bands and a concentration of WGA-reactive signal into two markedly more intense bands (Fig. 1D).

**Figure 1.**
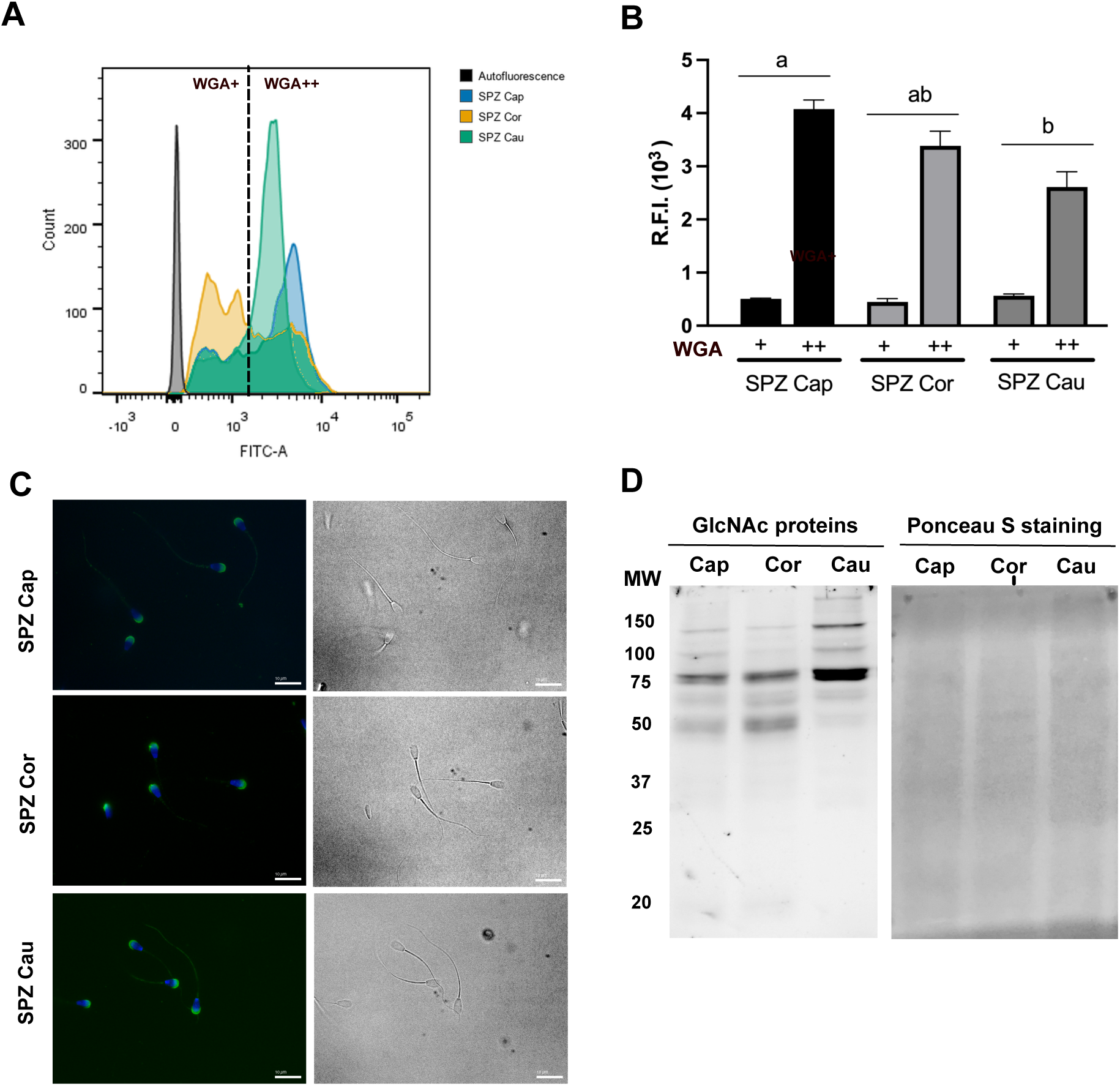
Changes in *N*-acetylglucosamine (GlcNAc) residues on the bovine sperm surface during epididymal transit. (A) Spermatozoa (SPZ) obtained from the epididymal caput (Cap), corpus (Cor), and cauda (Cau) were incubated with WGA-FITC lectin and analyzed by flow cytometry. (B) Relative fluorescence intensity (RFI) of each sperm population was quantified. The (+) population corresponds to sperm with lower RFI, and the (++) population to sperm with higher RFI. Data were analyzed by one-way ANOVA followed by Tukey’s post hoc test. Different letters indicate statistically significant differences (P < 0.05; mean ± SEM; n = 7 independent bulls). (C) Sperm from each epididymal region were stained with WGA-FITC and examined by fluorescence microscopy. (D) Patterns of GlcNAc-bearing proteins were analyzed in isolated sperm membrane fractions by lectin blot. Proteins were separated by SDS-PAGE, transferred to nitrocellulose membranes, and detected with biotin-WGA. Ponceau staining was used as a loading control.

### 3.2. **β**-*N*-acetylglucosaminidase (**β**-NAG) from cauda epididymal fluid can modify GlcNAc content on bovine spermatozoa

We previously demonstrated that β-NAG activity increases significantly toward the cauda epididymis in sexually mature bulls [10], suggesting that this luminal glycosidase is a strong candidate for driving the regional GlcNAc remodeling observed during epididymal transit. Building on this, we next evaluated whether β-NAG is directly responsible for GlcNAc modification on immature spermatozoa. Incubation of caput spermatozoa with cauda fluid under optimal conditions for enzyme activity resulted in a significant reduction in WGA–FITC labeling (P < 0.05), an effect that was partially reversed by co-incubation with the selective β-NAG inhibitor VP150, confirming the specificity of this enzymatic activity (Fig. 2A–B). These findings demonstrate that β-NAG in epididymal fluid is enzymatically active and may contribute to GlcNAc remodeling during sperm maturation.

**Figure 2.**
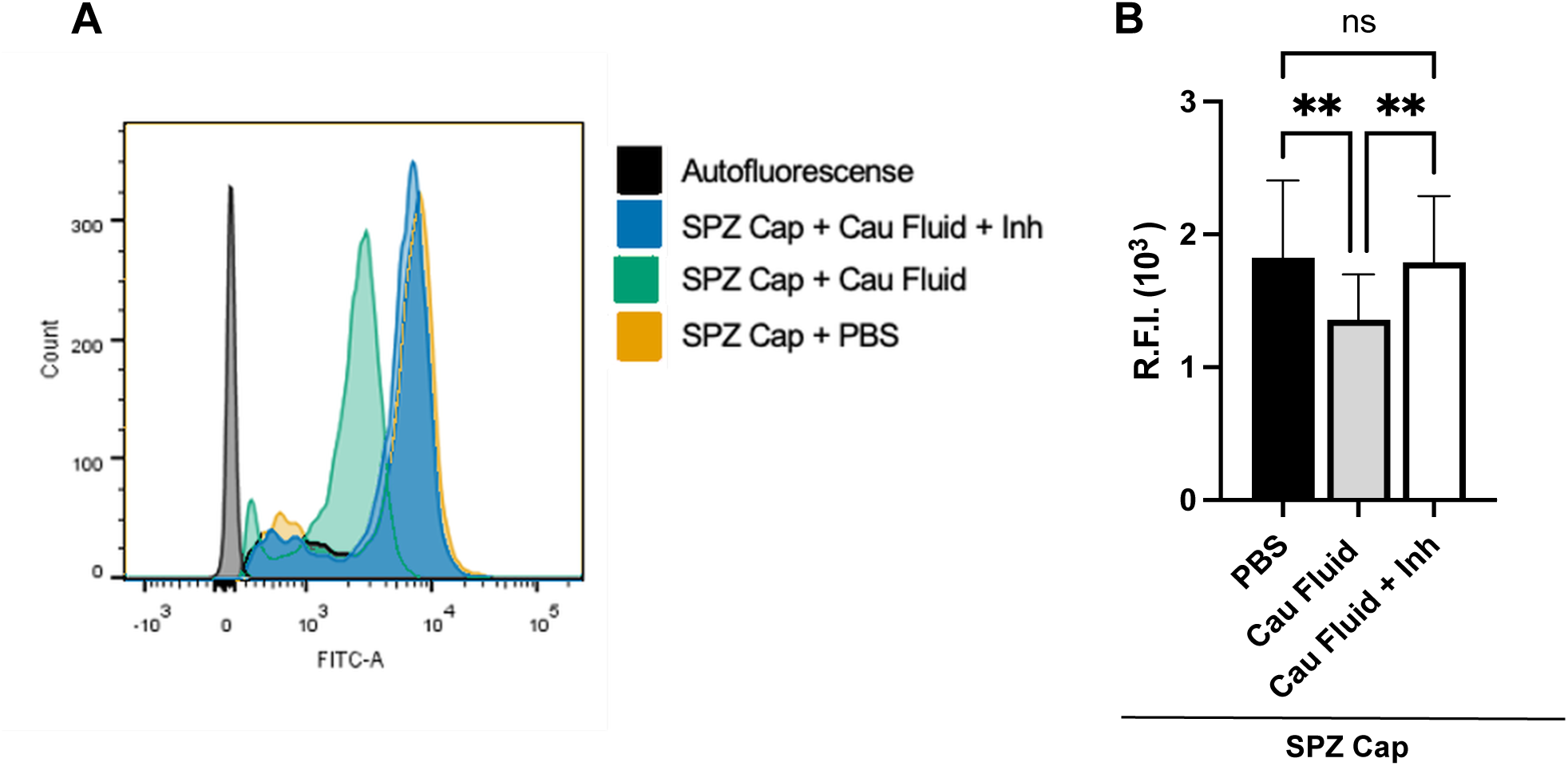
*N*-acetyl-β-D-glucosaminidase (β-NAG) from cauda epididymis fluid modifies the GlcNAc content on epididymal caput spermatozoa. (A) Spermatozoa (SPZ) obtained from caput epididymis were incubated with cauda fluid either in the absence or in the presence of the β-NAG specific inhibitor VP150 and the GlcNAc content was analyzed by flow cytometry. (B) Data were analysed by one-way ANOVA followed by a Tukey’s multiple comparisons test. (**) Significantly different from all the conditions used (P<0.05; mean ± SEM; n =14 independent bulls).

### 3.3. Capacitation and the acrosome reaction modulate surface GlcNAc in ejaculated spermatozoa

To assess whether surface GlcNAc levels are altered during *in vitro* capacitation and the acrosome reaction, ejaculated spermatozoa were exposed to the specific conditions for each process. Ejaculated spermatozoa exhibited a high proportion of WGA^++^ cells under basal conditions. Incubation in capacitating medium elicited modest changes, whereas calcium ionophore treatment significantly decreased the proportion of WGA^++^ cells (P < 0.05; Fig. 3A–B). Fluorescence microscopy confirmed a reduction in acrosomal labeling following acrosome reaction induction (Fig. 3C). Lectin blot analysis revealed changes in GlcNAc-bearing membrane proteins associated with these physiological processes (Fig. 3D).

**Figure 3.**
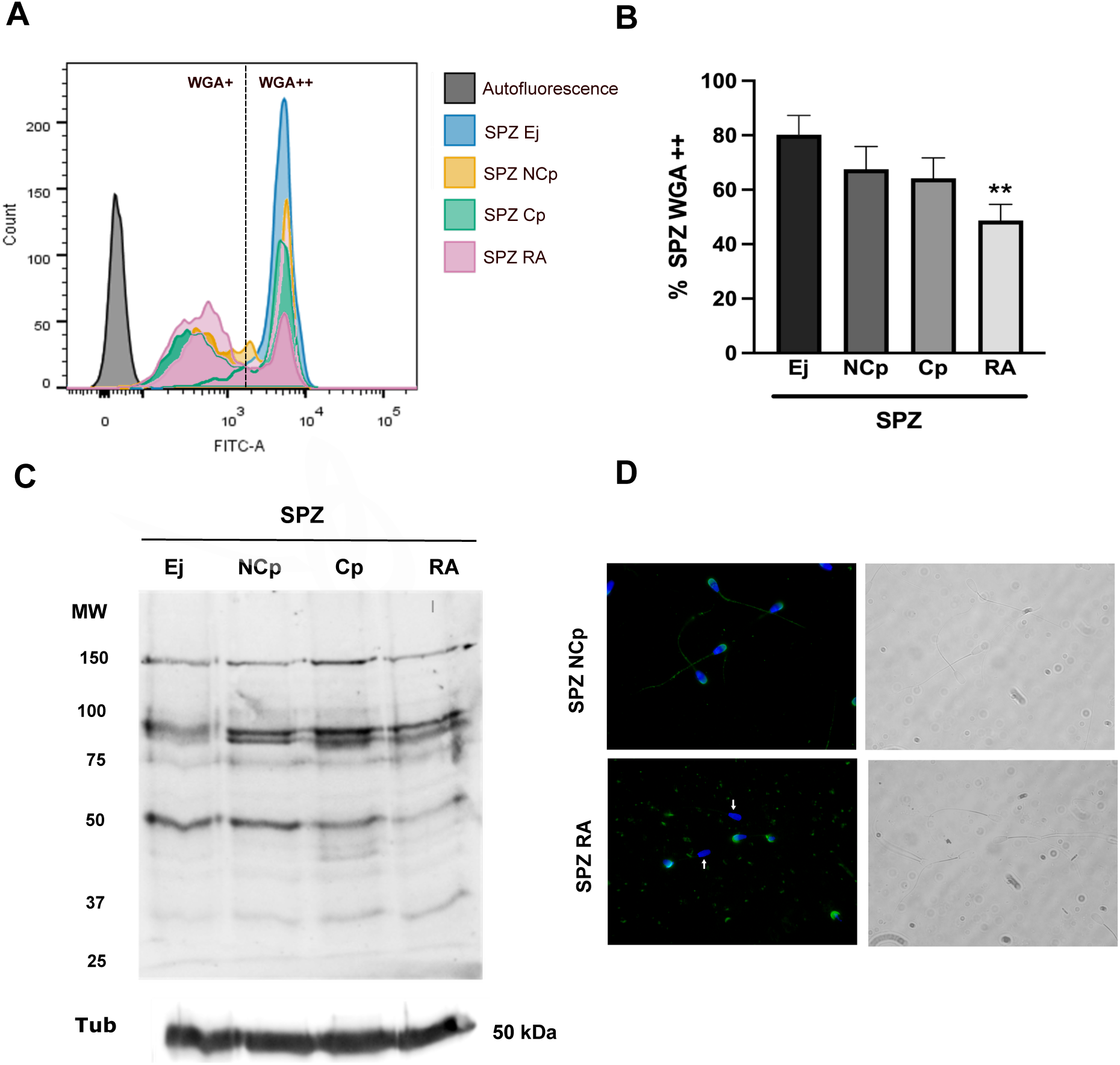
Changes in *N*-acetylglucosamine (GlcNAc) content associated with capacitation and the acrosome reaction in bovine ejaculated spermatozoa (SPZ). (A) Ejaculated bovine spermatozoa were thawed (SPZ Ej) and incubated under different conditions: non-capacitating medium (SPZ NCp), capacitating medium (SPZ Cp), or with calcium ionophore to induce the acrosome reaction (SPZ RA). After treatments, sperm were labeled with FITC-WGA lectin and analyzed by flow cytometry to quantify surface GlcNAc. (B) The percentage of sperm within the high-fluorescence population (FITC++) was quantified for each condition. (**) Significantly different from RA (P < 0.05, one-way ANOVA followed by Tukey’s post-hoc test; mean ± SEM; n = 5). (C) Patterns of GlcNAc-bearing proteins were assessed in SPZ membrane preparations by lectin blot using biotin-WGA on proteins electrotransferred to nitrocellulose membranes. Alpha-tubulin was used as loading control. (D) Representative fluorescence microscopy images of FITC-WGA-labeled spermatozoa under each condition. White arrows indicate WGA-negative spermatozoa.

### 3.4. Spermatozoa that bind to oviductal epithelial cells (BOECs) display GlcNAc residues on their surface

To investigate whether surface GlcNAc is associated with the ability to bind to the oviductal epithelium, ejaculated spermatozoa were co-incubated with BOEC monolayers. Microscopy analysis revealed that most spermatozoa attached to BOECs were WGA+ (Fig. 4A). Quantification confirmed that the proportion of WGA+ spermatozoa bound to BOECs was significantly higher than that of WGA spermatozoa (P < 0.001; Fig. 4B). Non-adherent spermatozoa were subsequently analyzed by flow cytometry to evaluate both membrane integrity and the presence of surface GlcNAc residues. Dual staining with propidium iodide (PI) and WGA revealed that a higher proportion of non-bound spermatozoa were PI+, indicating compromised membrane integrity. Notably, despite this increase in non-viable cells, a substantial fraction of these spermatozoa retained WGA binding, demonstrating that the presence of GlcNAc residues on the sperm surface is not restricted to viable cells (Fig. 4C–D). Additionally, preincubation of spermatozoa with WGA significantly reduced sperm adhesion (P < 0.05; Fig. 4E), supporting a role for GlcNAc residues in SPZ–BOEC interactions.

**Figure 4.**
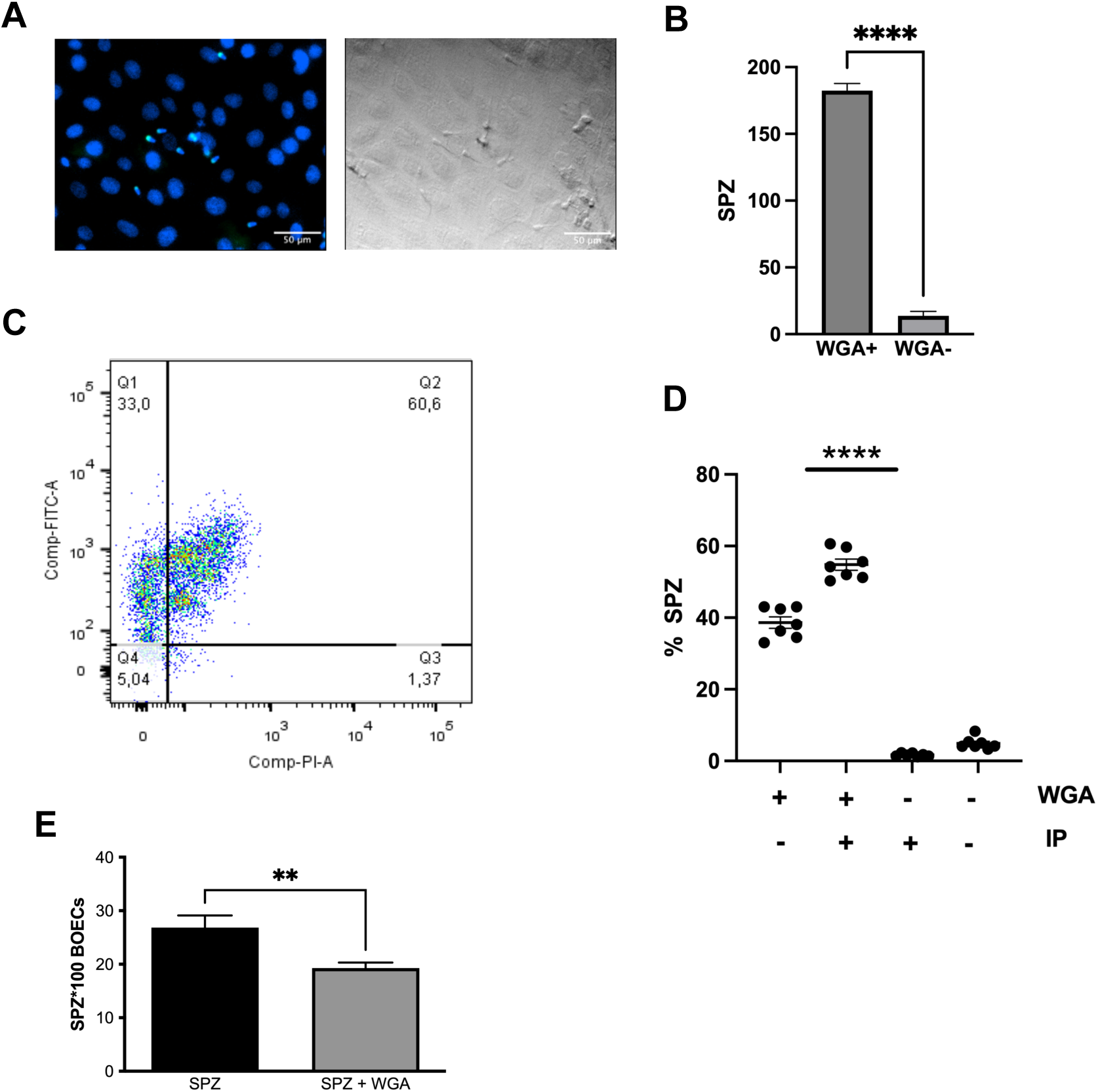
*N*-acetylglucosamine (GlcNAc) is present on the surface of spermatozoa that bind to BOECs. (A) Spermatozoa (SPZ) after co-incubation with BOECs, stained with Hoechst and WGA, visualized by fluorescence microscopy and bright field. (B) Quantification of WGA spermatozoa attached to BOECs. A total of 200 cells were quantified across 5 different samples. (****) Significant difference between WGA and WGA spermatozoa attached to BOECs, compared to control (t-test, p < 0.001). (C) Following co-culture, unbound spermatozoa were evaluated for surface GlcNAc content and viability using propidium iodide (PI) by flow cytometry. (D) Double-staining data were quantified in quartiles (Q1–Q4) across 4 different samples. (****) Significantly different from all other combinations. **(E)** BOEC monolayers were pre-treated with WGA prior to co-incubation with ejaculated spermatozoa. The number of spermatozoa bound to BOECs was quantified by fluorescence microscopy following nuclear staining with Hoechst. Bars represent the mean number of SPZ bound per 100 BOECs ± SEM. (N=4) (**) Significant difference compared to control (p < 0.05).

### 3.5. Complementary analyses

#### 3.5.1. The inhibitor VP150 effectively reduces β-NAG activity in cauda epididymal fluid

The molecular structure of the β-NAG inhibitor VP150 is shown in Supplementary Figure 1A. Measurement of enzymatic activity in cauda epididymal fluid from four bulls demonstrated a significant decrease in β-NAG activity in the presence of VP150, confirming its selectivity and efficacy under the experimental conditions used (Suppl. Fig. 1B).

#### 3.5.2. BOEC characterization and sperm viability during adhesion assays

The epithelial identity of BOEC cultures was confirmed by immunofluorescence for cytokeratin which showed a significantly greater proportion of Cyt cells relative to Vim cells (P < 0.01; Suppl. Fig. 2 A–B). During co-culture assays, spermatozoa bound to BOECs retained membrane integrity as verified by PI staining, which showed a predominant proportion of viable (PI) attached spermatozoa (Suppl. Fig. 3A–B).

## 4. DISCUSSION

Carbohydrate moieties on glycoproteins are essential mediators of cell–cell recognition, and their remodeling on the sperm surface is a hallmark of male gamete maturation and function [29]. The present study provides a systematic characterization of surface N-acetylglucosamine (GlcNAc) dynamics in bovine spermatozoa, linking epididymal maturation, functional activation, and sperm-oviduct epithelial adhesion. Using complementary lectin-based approaches, including flow cytometry, fluorescence microscopy, and lectin blot analysis, we demonstrate that the progressive change in surface GlcNAc content during epididymal transit is not simply descriptive but is causally linked to luminal β-*N*-acetylglucosaminidase (β-NAG) activity, which actively remodels GlcNAc residues on immature spermatozoa.This is consistent with our previous report of a significant increase in β-NAG activity toward the cauda in reproductively mature bulls compared with bulls that had not yet reached reproductive maturity [10], further supporting a role for this enzyme in sperm maturation.

We further show that capacitation and the acrosome reaction differentially modulate surface GlcNAc, and that spermatozoa bearing surface GlcNAc are preferentially recruited to bovine oviductal epithelial cell (BOEC) monolayers. Collectively, these findings advance our understanding of the glycobiological determinants that regulate sperm function in cattle.

Our flow cytometric data revealed progressive modifications in the proportion of WGA^++^ spermatozoa from the caput to the cauda epididymis, accompanied by changes in relative GlcNAc-bearing protein profiles by lectin blot. These results are consistent with earlier reports of epididymis-dependent glycocalyx remodeling in mammals [1,8,18,21,30] and extend them by providing quantitative regional data in the bovine model. Importantly, the spatial distribution of GlcNAc, as assessed by fluorescence microscopy, remained predominantly acrosomal throughout epididymal transit, suggesting that regional enrichment occurs mainly within a restricted and functionally relevant membrane domain.

Among these earlier reports, a recent study using a broad panel of lectins similarly described extensive glycocalyx remodeling in bull spermatozoa during epididymal transit [18]. However, the specific WGA-reactive protein pattern we observed diverges from this previous report: whereas that study described the emergence of additional WGA-reactive bands toward the cauda, our lectin blot analysis revealed the opposite trend: a progressive loss of GlcNAc-bearing bands accompanied by the concentration of WGA reactivity into two markedly more intense bands. This divergence may be explained, at least in part, by differences in membrane isolation methodology: whereas that study purified plasma membranes by discontinuous sucrose density-gradient ultracentrifugation, our protocol relied on hypo-osmotic lysis followed by differential centrifugation, which may yield membrane fractions of different purity and subcellular composition. Alternatively, or in addition, this discrepancy may reflect a genuine biological difference in the fate of GlcNAc-bearing glycoproteins: rather than a generalized increase in glycan and glycoprotein complexity during epididymal transit, specific glycoprotein subsets recognized by WGA may instead undergo selective enzymatic trimming, potentially mediated by luminal β-NAG activity, resulting in the simplified and concentrated banding pattern observed toward the cauda.

The epididymis secretes an array of glycosidases, glycosyltransferases, and lectins into its luminal fluid, constituting a microenvironment capable of extensively modifying the sperm glycocalyx [6]. In this context, the progressive modification in WGA labeling from caput to cauda is likely to reflect both the addition of GlcNAc-terminal glycans and the unmasking of pre-existing residues by the removal of sialyl caps or other terminal sugars. This shift was further associated with a change in population structure: while caput and corpus spermatozoa comprised a mixture of subpopulations displaying distinct WGA labeling patterns, the sperm population became markedly more homogeneous toward the cauda, as revealed by flow cytometry. Notably, our data also showed distinct GlcNAc-bearing protein profiles across epididymal regions by lectin blot, consistent with the concept of molecular remodeling during sperm maturation reported in other species [1,31]. Whether these changes reflect *de novo* glycosylation via epididymosome-mediated transfer or enzymatic modification of existing glycans warrants further investigation.

In this context, to explore enzymatic mechanisms, we incubated caput spermatozoa with cauda epididymal fluid and observed a significant reduction in WGA-FITC labeling, an effect that was partially reversed by co-incubation with the selective β-NAG inhibitor VP150. These results provide direct evidence that β-NAG activity in epididymal luminal fluid is enzymatically active and can modify GlcNAc residues on immature spermatozoa. This finding is consistent with previous reports from our group showing that β-NAG activity increases in the cauda epididymal fluid relative to the caput and corpus in sexually mature bulls [10], and supports a model in which luminal glycosidase activity contributes to epididymal GlcNAc remodeling.

The role of glycosidases in epididymal sperm maturation has long been proposed [30], yet direct experimental evidence linking specific enzymes to specific glycan changes on sperm has remained sparse. Our results with VP150 establish a causal link between β-NAG activity and GlcNAc reduction on caput spermatozoa, providing a functional correlate of the regional enzymatic activity data. Interestingly, given that the cauda environment generates mature cauda-like GlcNAc patterns, the fact that the net effect is a reduction suggests that β-NAG-mediated de-glycosylation may represent a terminal trimming step in GlcNAc remodeling, or that the enzyme acts selectively on immature glycoforms. Alternatively, the paradoxical relationship between regional GlcNAc increase during epididymal transit and the β-NAG-mediated removal of GlcNAc *in vitro* may reflect the complexity of *in vivo* glycan remodeling, where multiple competing enzymatic activities and transfer mechanisms operate simultaneously [1,6,31].

Furthermore, we investigated the dynamics of GlcNAc residues during capacitation and the acrosome reaction. We found that while incubation in capacitating medium did not introduce significant changes in WGA labeling, induction of the acrosome reaction with calcium ionophore A23187 produced a significant decrease in the proportion of WGA spermatozoa, accompanied by a marked reduction in acrosomal labeling by fluorescence microscopy. These results are consistent with the established role of acrosomal exocytosis in releasing or redistributing acrosome-associated glycoproteins [32] and extend prior observations of surface changes reported during capacitation in bovine [33] and dolphin [34] spermatozoa, and during the acrosome reaction in porcine [35] and human [36] spermatozoa.

During bovine capacitation, major membrane changes include cholesterol efflux, protein tyrosine phosphorylation, and lipid raft reorganization [32]. These events may reorganize rather than deplete surface glycans. In contrast, the acrosome reaction involves extensive membrane fusion between the plasma membrane and the outer acrosomal membrane, resulting in the release and loss of acrosomal glycoproteins bearing GlcNAc moieties from the plasma membrane [1]. Our lectin blot analysis of total protein extracts from ejaculated, capacitated, and acrosome-reacted spermatozoa corroborated this interpretation by revealing distinct GlcNAc-bearing protein profiles associated with each physiological state. The use of total protein extracts, rather than isolated membrane fractions, in this experiment was intentional, as it allowed full accounting of acrosomal proteins that would be lost or redistributed during exocytosis, thus providing a more complete view of glycoprotein dynamics.

Perhaps the most functionally significant finding of this study is the preferential adhesion of WGA spermatozoa to BOEC monolayers. Quantitative analysis confirmed a significantly higher proportion of WGA cells among bound spermatozoa compared with unbound counterparts, and preincubation of spermatozoa with unconjugated WGA significantly reduced BOEC adhesion. These results provide direct experimental evidence that surface-exposed GlcNAc residues participate in sperm-BOEC interactions, supporting a role for this glycan in the formation or stability of the bovine sperm reservoir. Interestingly, GlcNAc residues have gained additional relevance considering recent findings identifying specific transporters such as SLC35G3, responsible for UDP-GlcNAc transport, whose deficiency severely compromises the formation of glycoproteins essential for passage through the uterotubal junction and binding to the zona pellucida [11].

The oviduct isthmus is the principal site of sperm reservoir formation in cattle, and sperm-BOEC adhesion is mediated by carbohydrate-lectin interactions at the gamete-epithelial interface [13,37]. The present results suggest that WGA-reactive sperm surface residues may serve as complementary ligands for such oviductal carbohydrate-binding proteins. The competitive inhibition of sperm adhesion by pre-incubation with WGA further supports a lectin-mediated mechanism in which GlcNAc residues on the sperm surface engage with specific carbohydrate-binding partners on the oviductal epithelium. This interpretation is consistent with observations showing that specific glycan motifs on the oviductal surface, including glycosaminoglycans, regulate sperm adhesion to the isthmic reservoir [37,38].

Flow cytometric dual-staining with WGA-FITC and propidium iodide revealed that a substantial proportion of non-adherent spermatozoa were PI, indicating compromised membrane integrity in this subpopulation. Notably, even within the PI fraction, a significant proportion retained WGA binding, demonstrating that GlcNAc exposure is not exclusively a property of viable cells. This observation may be relevant from a selective perspective: if BOEC binding favors WGA /PI spermatozoa, as suggested by the viability assessment during co-culture, the oviductal reservoir would preferentially retain membrane-intact cells expressing surface GlcNAc, consistent with a selective function of the reservoir in promoting the fertilizing capacity of the sperm population [29,37].

This study provides an integrative view of GlcNAc dynamics from epididymal maturation through sperm-oviduct interaction in the bovine. The findings are particularly relevant in the context of assisted reproduction technologies (ART), where frozen-thawed semen, as used here, can undergo cryopreservation-associated changes in surface glycan composition [1,39]. Understanding the glycobiological requirements for successful sperm-oviduct adhesion may inform strategies to improve sperm selection and *in vitro* fertilization outcomes in cattle, a species of significant economic importance. Future studies should focus on identifying the specific GlcNAc-bearing glycoproteins involved in BOEC adhesion, using proteomic and glycoproteomic approaches. The identity of the complementary lectin(s) on the oviductal epithelium that recognize sperm GlcNAc, as well as the signaling events triggered by this interaction, also remain to be elucidated.

In summary, this study demonstrates that surface GlcNAc residues undergo dynamic regulation at multiple stages of bovine sperm biology: they change progressively during epididymal maturation through mechanisms that include luminal β-NAG enzymatic activity, are substantially redistributed and lost following the acrosome reaction, and are associated with the preferential adhesion of spermatozoa to oviductal epithelial cells. These findings identify surface GlcNAc as a functionally relevant glycan at the sperm-oviduct interface and provide a mechanistic foundation for future studies on glycocalyx-mediated sperm selection in the bovine reproductive tract.

## Author Contributions

Conceptualization, P.A.A. and A.C.A.; Methodology, P.A.A., F.C. and N.L.L.; Visualization, P.A.A. and A.C.A.; Formal analysis, P.A.A. F.L. and N.L.L.; Investigation, P.A.A. and A.C.A.; Resources, I.R., A.C.A. and M.A.S.; Writing – Original Draft, A.C.A. and P.A.A.; Writing – Review & Editing, A.C.A. and M.A.S.; Supervision, A.C.A.; Funding Acquisition, M.A.S. and A.C.A. All authors have read and agreed to the published version of the manuscript.

## Funding

This research was funded by Agencia Nacional de Promoción Científica y Tecnológica (ANPCyT), grant PICT-2018-01286; by Consejo Nacional de Investigaciones Científicas y Técnicas (CONICET), grant PIP 11220200103226CO01; and by Secretaría de Investigación, Internacionales y Posgrado (SIIP), Universidad Nacional de Cuyo, grant M002-T1.

## Ethics Statement

This study did not involve live-animal experimentation. No animals were sacrificed specifically for this study; bovine reproductive tracts (epididymides and oviducts) were obtained as by-products of routine commercial slaughter, independently of and unrelated to this research, at a SENASA-licensed slaughterhouse (María del Carmen, Corralitos, Mendoza, Argentina), operating under the Argentine Regulation for Inspection of Animal Products, Sub-products and Derivatives (Decreto N° 4238/1968) and the animal welfare requirements for livestock slaughter established by SENASA Resolution 1697/2019 (as amended). As no live animals were manipulated for the purposes of this work, ethical review under CONICET Resolution 1047/05 - which governs live-animal research protocols evaluated by Institutional Committees for the Care and Use of Laboratory Animals (CICUAL) - was not aplicable. The ARRIVE 2.0 guidelines were not applicable, as this study did not involve procedures on living animals.

## Data Availability Statement

The data supporting the findings of this study are available from the corresponding author upon reasonable request.

## Acknowledgments

The authors thank Ms. Andrea Lafalla Manzano and Mr. Gabriel Houri (Flow Cytometry Facility, Facultad de Ciencias Médicas, Universidad Nacional de Cuyo) for their valuable technical assistance and consultation during flow cytometry experiments. The authors also thank María del Carmen slaughterhouse, and particularly D.V.M. Juan Ponzi and D.M.V. Avelino Maure, for their assistance and technical support in sample collection.

## Conflicts of Interest

The authors declare no conflict of interest.

---

**Supplementary Figure 1.**
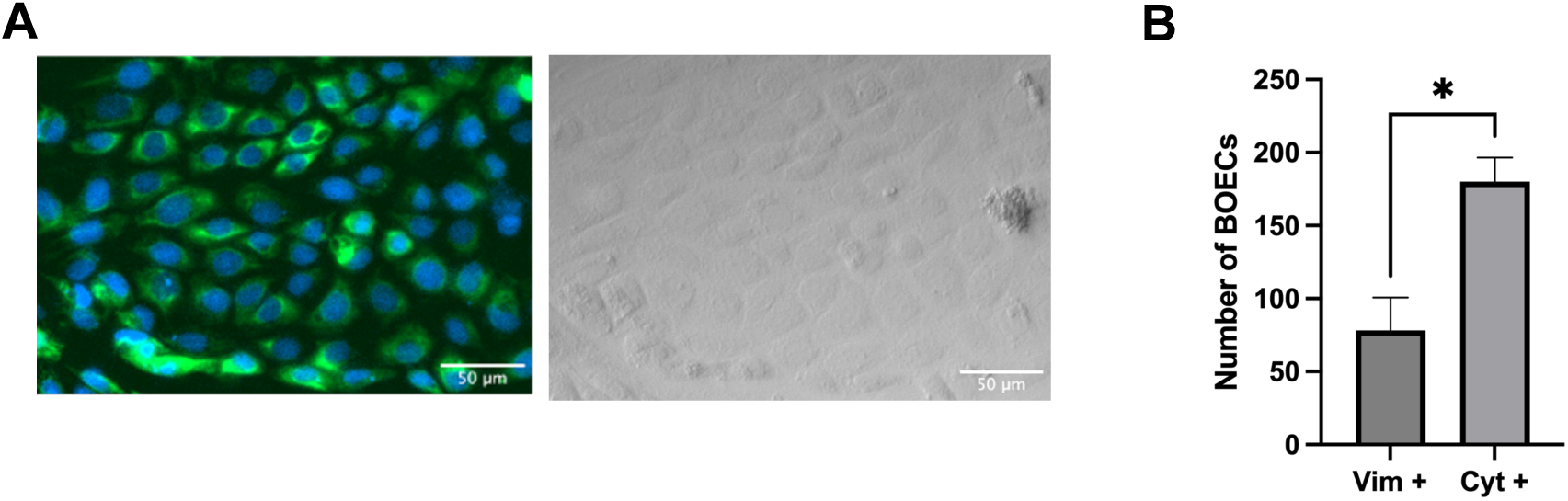
Inhibition of β-*N*-acetylglucosaminidase (β-NAG) activity in bovine cauda epididymal fluid by the VP150 inhibitor. (A) Molecular structure of the β-NAG inhibitor VP150 used in this study. (B) Enzymatic activity of β-NAG measured in cauda epididymal fluid from four independent bulls, expressed as mean ± SD. The basal activity of β-NAG in untreated samples was compared with the activity measured in the presence of VP150. A significant reduction in β-NAG activity was observed when the enzyme was incubated with the inhibitor, confirming the inhibitory efficacy of VP150 under the assay conditions.

**Supplementary Figure 2.**
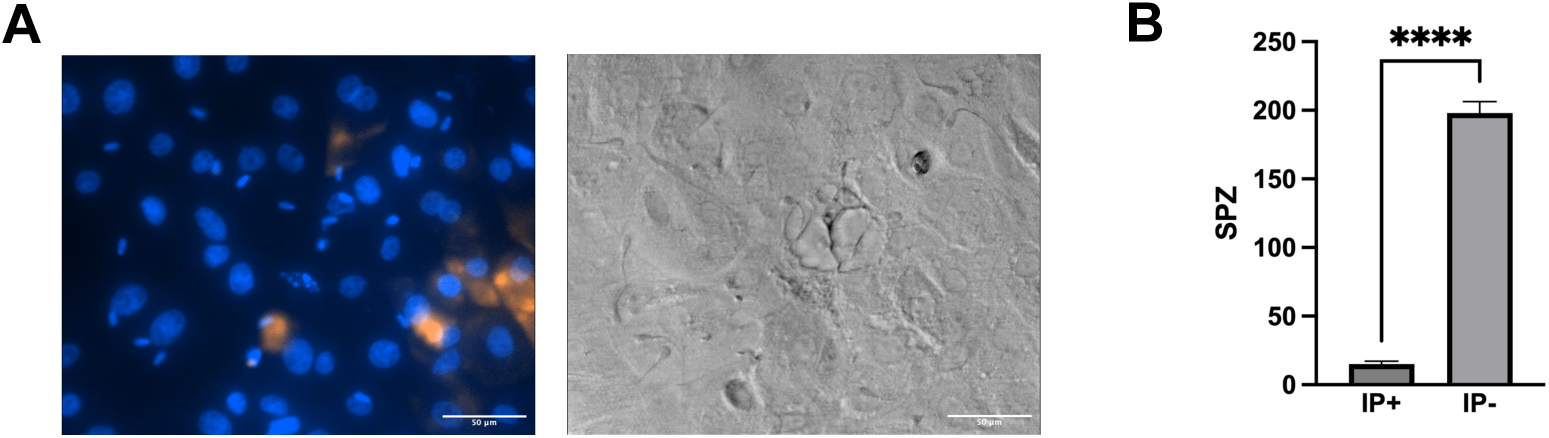
Phenotypic characterization of bovine oviductal epithelial cells (BOECs). (A) Immunofluorescence microscopy of BOECs cultured to confluence. Cells were stained with anti-cytokeratin antibody followed by a FITC-conjugated secondary antibody. (B). The number of vimentin-positive (Vim) and cytokeratin-positive (Cyt) cells was quantified from the immunofluorescence images. A total of 250 cells were counted in simultaneous preparations, one stained for Vim and the other for Cyt, followed by an anti-rabbit FITC-conjugated secondary antibody (N=3). (*) Significant difference between the two populations (t-test; p < 0.05).

**Supplementary Figure 3.**
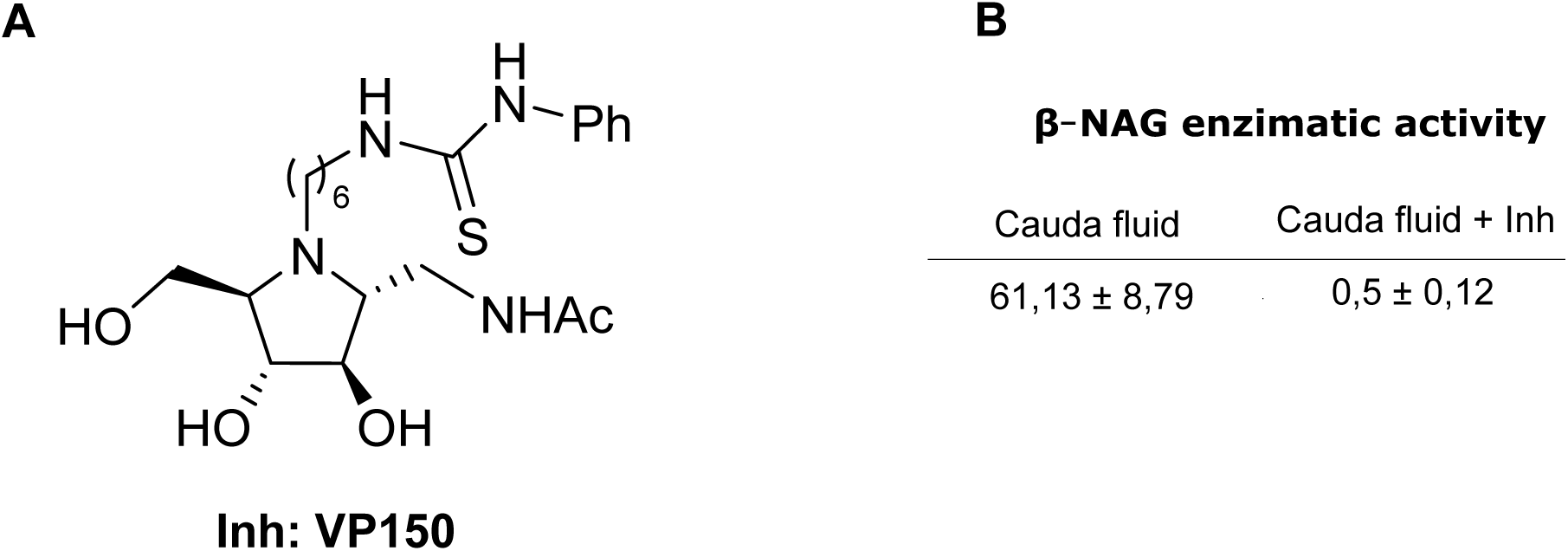
Evaluation of sperm viability during binding to BOECs using propidium iodide (PI). (A) Fluorescence microscopy of BOECs co-incubated with spermatozoa and stained with propidium iodide (PI) to assess membrane integrity. The corresponding bright-field image is shown for morphological reference. (B) Quantification of PI-positive (PI; non-viable) and PI-negative (PI; viable) spermatozoa bound to BOECs. A paired t-test revealed a significant difference between both populations (**** p < 0.001).

## Notes

### Competing Interest Statement

The authors have declared no competing interest.

### Summary of Updates

The Introduction and Discussion sections were updated to improve the comparison of the new results with the existing literature. Additionally, Figures 1 and 3 were revised and updated.

